# Assessing the robustness of deep learning based brain age prediction models across multiple EEG datasets

**DOI:** 10.1101/2025.05.20.655022

**Authors:** Thomas Tveitstøl, Mats Tveter, Christoffer Hatlestad-Hall, Hugo L Hammer, Denis A Engemann, Ira R J Hebold Haraldsen

## Abstract

The increasing availability of large electroencephalography (EEG) datasets enhances the potential clinical utility of deep learning (DL) for cognitive and pathological decoding. However, dataset shifts due to variations in the population and acquisition hardware can considerably degrade the model performance. We systematically investigated the generalisation of DL models to unseen datasets with different characteristics, using age as the target variable. Five datasets were used in two different experimental setups, including (1) leave-one-dataset-out (LODO) and (2) leave-one-dataset-in (LODI) cross validation. A comprehensive set of 1805 different hyperparameter configurations was tested, including variations in the DL architectures and data pre-processing. The performance varied across source/target dataset pair. Using LODO, we obtained Pearson’s r values of {0.63, 0.84, 0.75, 0.23, 0.10} and *R*^2^ values of {-0.01, 0.63, 0.41, −4.66, −70.98}. For LODI, the results varied in Pearson’s r from −0.11 to 0.84 and *R*^2^ values from −704.89 to 0.65, depending on the source and target dataset. Adjusting the model intercepts using the average age of the target dataset substantially improved some *R*^2^ scores. Our results show that DL models can learn age-related EEG patterns which generalise with strong correlations to datasets with broad age spans. The most important hyperparameter was to use the frequency range between 1 and 45Hz, rather than a single frequency band. The second most important hyperparameter effect depended on the experimental setup. Our findings highlight the challenges of dataset shifts in EEG-based DL models and establish a benchmark for future studies aiming to improve the robustness of DL models across diverse datasets.

## 1. Introduction

Artificial intelligence (AI) is becoming an increasingly powerful and impactful tool in health and medicine [1, 2]. The AI-Mind project (www.ai-mind.eu) aims to develop prognostic AI-based tools for dementia risk assessment in persons with mild cognitive impairment (MCI) [3]. A key modality in this project is to leverage high-density electroencephalography (EEG), which directly reflects the electrical activity of the brain, to predict the risk of dementia. Deep learning (DL) directly applied on the EEG signals is of interest, due to its ability to learn complex patterns beyond hand-crafted features. Instead, DL processes the data in multiple layers with different levels of abstraction [4]. With more and larger datasets becoming available, DL becomes increasingly relevant, as these models are effective at handling large amounts of unstructured data for successful pattern recognition. However, for generalised use of a model, training AI models on only a single dataset may be insufficient, even with large sample sizes [5]. Dataset shifts are known difficulties, where systematic differences between the training data and new data from different sites can considerably degrade model performance [6, 7]. Although the cross-dataset setting has been the focus in some studies, these works are mostly limited to brain computer interface applications [8, 9, 10, 11, 12] and emotion recognition [13, 14, 15, 16, 17]. Furthermore, many of these studies focus on domain adaptation and transfer learning [14, 18, 15, 19, 9, 10, 20, 11, 17, 12], which typically requires data from the target domain. However, there is currently a lack of studies which focus on resting-state EEG and domain generalisation, where datasets with different characteristics are fully excluded from the model training. Investigating this is therefore of high importance, as ensuring that EEG-based DL models generalise well across different clinical sites, populations, and hardware variations, is crucial for clinical application.

To investigate the ability of DL models to extract patterns that generalise across datasets, age was selected as the target variable. Brain aging is usually associated with neuronal apoptosis and a loss of synaptic contacts [21], and is the main risk factor for neurodegenerative disorders, including Parkinson’s disease, amyotrophic lateral sclerosis, stroke, and Alzheimer’s disease [21, 22]. With machine learning, patterns of both healthy and pathological aging can be extracted in a data driven manner. Previous studies using MRI have shown an association between the difference between chronological and predicted brain age, and pathological conditions [23, 24]. Although these findings are from studies using MRI data, which reflects different underlying biological mechanisms, EEG has indeed been used to study brain aging [25, 26], and was recently benchmarked for more specific predictive modeling [27, 28]. Importantly, [28] highlighted that choices of EEG processing can substantially impact prediction performance. The selection of age as target variable also allows the use of multiple open-source datasets, as it is commonly available in EEG datasets. Furthermore, due to the possible relevance of brain age in cognitive health, we argue that assessing the ability of DL models to generalise to unseen datasets for predicting brain age may be an important step towards assessing the ability of DL models to extract clinically and pathologically relevant EEG features that generalise to new datasets.

In this study, we investigated the cross-dataset generalisation capabilities of DL models using five resting-state EEG datasets with different characteristics. An extensive set of experiments with randomly sampled hyperparameters (HPs) and design choices was carried out. This served mainly three purposes: (1) to test a wide variety of HP configurations (HPCs) and assess their impact on cross-dataset generalisation, (2) to avoid reliance on a single set of HPCs, and (3) to document the key aspects of our model development and model selection. Our experiments may be categorised into leave-one-dataset-out (LODO) cross validation and leave-one-dataset-in (LODI) cross validation. The former is a cross validation strategy which trains on all but one dataset, while the latter trains on a single dataset and leaves the remaining ones for testing. With these cross validation variations, our aim was to cover the scenarios where multiple datasets (LODO) and a single dataset (LODI) are available for training.

## 2. Methods

To investigate the ability of DL models to generalise across EEG datasets, multiple datasets with different characteristics were combined. The experiments are categorised in two: (1) LODO and (2) LODI cross validation. In the former, cross validation was performed by leaving out a single dataset as the unseen test set. In the latter, cross validation was performed using only a single dataset for training, with the remaining datasets used for testing.

To avoid reliance on a single set of HPCs and design choices, we repeated the experiments for many sets of HPCs. See Figure 1 for a shallow overview. Our experiments varied in preprocessing and feature extraction steps, including EEG input duration, algorithm for channel repair and epoch removal, band-pass filtering, sampling frequency, and data normalisation. Furthermore, three DL architectures with randomly sampled HPs were used. To address dataset shifts, two domain adaptation techniques were used. This included domain adversarial learning [29] and Convolutional Monge Mapping Normalisation (CMMN) [30], which can also be adapted at test time. A strategy for sample re-weighting the loss function was implemented, whose impact ranged from equal weighting of all subjects to equal weighting of all datasets. Furthermore, different strategies were used for handling varied electrode configurations across the datasets. Finally, the HPs of the Adam [31] optimiser were randomly sampled. All code is publicly available at https://github.com/thomastveitstol/ CrossDatasetLearningEEG

**Figure 1.**
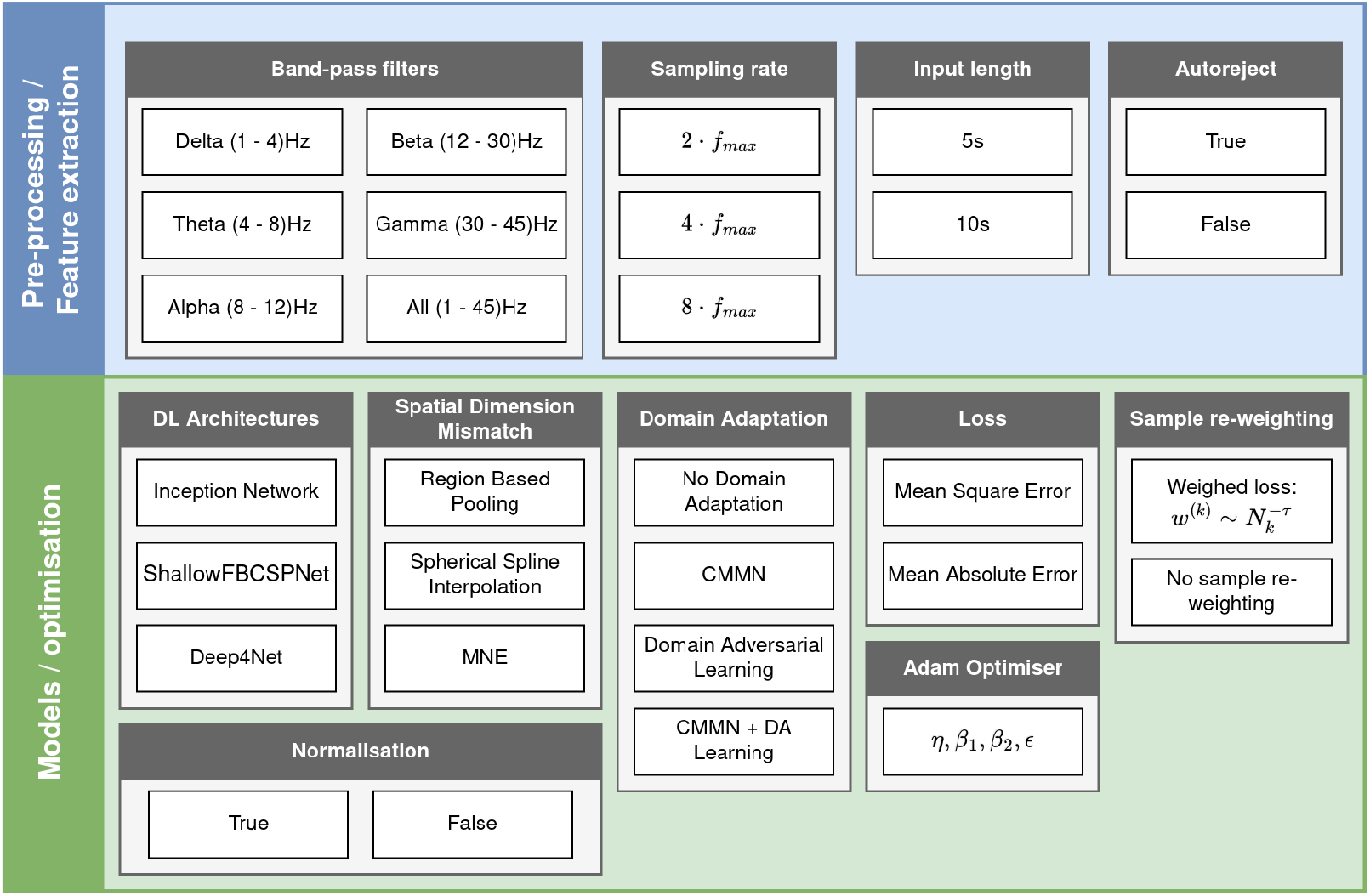
A shallow overview of the different hyperparameters. The hyperparameters of the DL architectures, region based pooling, CMMN, and domain adversarial learning, have been omitted from this figure. Abbreviations: DA (domain adversarial), CMMN (convolutional Monge mapping normalisation)

The HPs were randomly sampled from the distributions detailed in this section.

The datasets are also described in this section.

### 2.1. Data

The EEG data used for this study was limited to the resting state condition, where the participants had their eyes closed. In this subsection, the datasets are briefly described. For more comprehensive descriptions, see the original work (see below). Furthermore, the pre-processing and feature extraction steps are described.

#### 2.1.1. Datasets Figure 2 shows the age distributions of the different datasets used in this study

*TDBRAIN* [32] (www.brainclinics.com/resources) is a dataset from the Netherlands which contains N=1273 subjects with various psychiatric disorders. These psychiatric disorders include major depressive disorder (MDD; N=426), attention deficit hyperactivity disorder (ADHD; N=271), subjective memory complaints (SMC; N=119) and obsessive-compulsive disorder (OCD; N=75). The recordings were made with 26 channels using a Compumedics Quickcap or ANT-Neuro Waveguard Cap with sintered Ag/AgCl electrodes, based on the 10–10 electrode international system.

**Figure 2.**
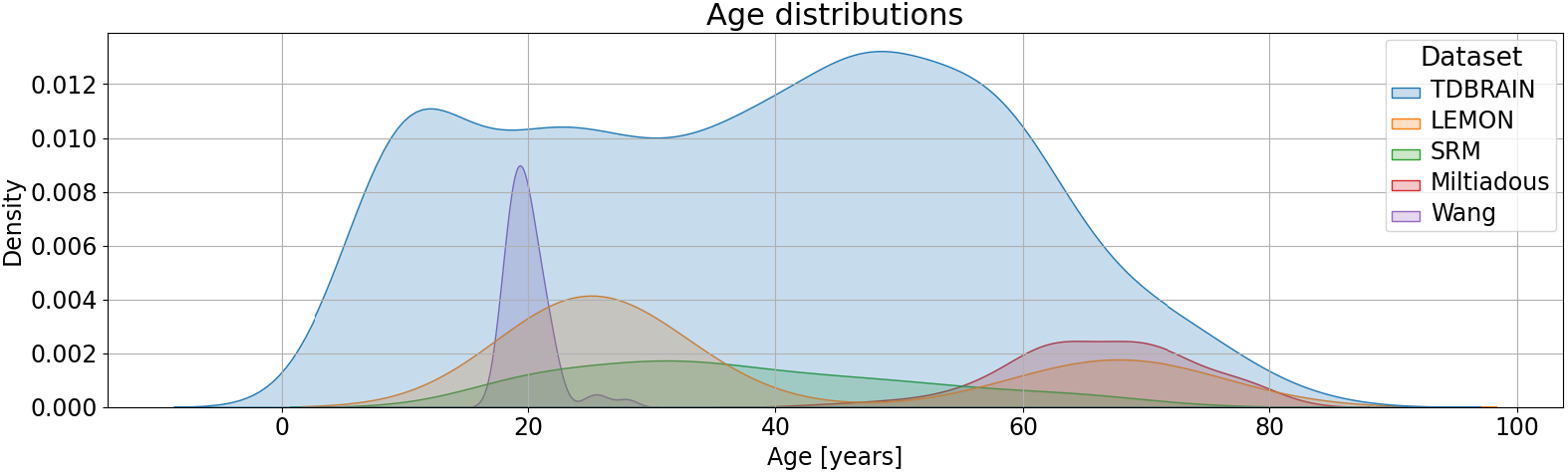
Age distributions of the different datasets.

*LEMON.* The LEMON dataset [33] used in this work consists of EEG from 203 healthy participants collected in Leipzig, Germany. It was recorded using 62-channel (61 scalp electrodes and one eye electrode) active ActiCAP electrodes following the extended international 10-20 system. Only the scalp electrodes were used, and missing channels were interpolated using the MNE method from MNE-Python [34]. The age intervals provided for the participants were averaged, e.g., a participant aged between 20 and 25 years was assigned an age of 22.5 years.

*SRM* (Sensory Response Modulation) [35] is a dataset with N=111 healthy controls, collected in Oslo, Norway. The EEG data was recorded during an SRM experiment and a resting-state session with a 64-channel (Ag-AgCl electrodes) BioSemi ActiveTwo system (BioSemi B.V., Amsterdam), following the electrode positional scheme of the extended 10-20 system. Only the resting-state data were used here.

*Miltiadous A dataset of EEG recordings from: Alzheimer’s disease, Frontotemporal dementia and Healthy subjects* [36] contains a total of 88 subjects, collected in Thessaloniki, Greece. Notably, it contains subjects with Alzheimer’s disease (N=36), frontotemporal dementia (N=23), and healthy controls (N=29). The recordings were made using a clinical EEG device (Nihon Kohden 2100) with 19 electrodes placed according to the international 10-20 system. For practical reasons, we use the name of the first author to refer to this dataset throughout the paper.

*Wang A test-retest resting and cognitive state EEG dataset* [37], contains N=60 subjects, collected at Southwest University in Chongqing, China. The EEG was recorded with either 63 or 64 Ag/AgCl active electrodes (the dataset available at OpenNeuro contains 62 reconstructed channels), with channel placement according to the extended international 10-20 system. For practical reasons, we use the name of the first author to refer to this dataset throughout the paper.

#### 2.1.2. Pre-processing and feature extraction

##### Input data

All pre-processing and feature extraction steps of the EEG data were implemented using MNE-Python [34] and autoreject [38]. The first 30 seconds of all EEGs were removed, as artifacts are more common in the first part of the EEG. The data were then band-pass filtered between 1 and 45 Hz. The continuous data were then segmented into non-overlapping epochs. Both 5 and 10 second windows were used for our experiments. For a single experiment (either LODO or LODI), the length was selected by random sampling with equal probabilities. One version of the data was cleaned with *autoreject* [38], while another version did not use any algorithm for channel repair or epoch rejection. Before running autoreject, the EEG data was resampled to 180Hz to reduce the runtime. Five epochs per sample were used. The input to the DL models, however, was a single epoch, with data split on subject level for training and validation. That is, the input to the DL models was a single epoch, but the total number of EEGs used for training was increased by a factor of five due to the inclusion of multiple epochs per original sample. The data was filtered into six frequency bands: *δ* (1-4Hz), *θ* (4-8Hz), *α* (8-12)Hz, *β* (12-30Hz), *γ* (30-45Hz), and all frequency bands (1-45Hz). Three different sampling rates were used per band-pass filtered version, sampling rate ∈ {2*f*_*max*_, 4*f*_*max*_, 8*f*_*max*_}, initially sampled with equal probabilities. However, as some of the DL architectures had a minimum number of required input time steps, the sampling rate was doubled upon failed model initialisation, if the sampling frequency was not already at the maximum. Finally, the EEG data was re-referenced to average. In total, 72 different versions of the EEG data were stored, and sampled with equal probabilities for each experiment.

##### Target data

The target values were z-normalised such that the training data had a mean of zero and unit standard deviation. For computing metrics, an inverse scaling was applied to the DL model outputs.

### 2.2. DL architectures

Three different DL architectures were used, and the architecture selection was made by random sampling with equal probabilities per experiment. Here, we provide a brief description of the architectures as well as the distributions from which their HPs were sampled from. For a more detailed description of the architectures, see the original papers.

For each experiment, there was a 50% probability that the inputs were normalised such that each input channel/region representation had zero mean and unit standard deviation. The prediction made for a given subject was made by averaging the predictions from the five EEG epochs.

#### 2.2.1. Inception network

Inception network [39] was originally designed and tested on a broad range of multivariate time series classification problems. With original HPCs, Inception network is comprised of two residual blocks, each such block containing three Inception modules connected in series. Each Inception module performs convolution with varied kernel sizes, and concatenates the resulting feature maps. Skip connection with a shortcut layer connects subsequent residual blocks.

In this work, we sampled the depth and the number of convolutional filters per layer (consistent through all layers). Here, we refer to the depth of the Inception network as the total number of Inception modules. The depth *d* ∈ N was sampled from the distribution *d* = 3[3^*n*^], *n* ∼*U* [0, 3], where [·] denotes the rounding operator. This favors fairly small depths (although, the original uses only a depth equal to six), which is pragmatic due to shorter run-time, but still allows for deep architectures as well. The number of convolutional filters *f* ∈ ℕ was sampled from a log uniform distribution as *f* = [2^*n*^], *n* ∼*U* [3, 6].

#### 2.2.2. Deep4Net

Deep4Net [40] was originally designed for brain computer interface (BCI), and is implemented in the braindecode package [41]. It consists of four convolutional-max pooling blocks, where the first convolutional block contains a temporal and a spatial filter. The subsequent three blocks use convolution and max pooling, where the number of convolutional units increases with a factor of two per convolutional block.

To allow for a smaller number of time samples, padding was added to the last three convolutional layers. The kernel length of all convolutional blocks in the temporal dimension was sampled from the discrete uniform distribution *l* ∼ *U*_ℕ_ [5, 15]. The number of convolutional layers in the first block was sampled from a discrete uniform distribution, *f*_1_ ∼ *U*_ℕ_ [15, 35]. The number of filters for the subsequent convolutional blocks was calculated as the double of its prior, to maintain the same architectural structure as the original implementation. That is, the number of filters for the *i*-th convolutional block was calculated as *f*_*i*_ = 2*f*_*i*−1_, ∀*i* ∈ {2, 3, 4}. The drop out rate prior to the final layer was sampled uniformly between 0 and 0.5.

#### 2.2.3. ShallowFBCSPNet

ShallowFBCSPNet [40] is an architecture developed for BCI tasks, and available in braindecode [41]. It was inspired by the filterbank common spatial pattern [42, 43] pipeline. As in Deep4Net, the first two layers are separate temporal and spatial convolutional layers. This is followed by a squaring nonlinearity, mean pooling, and a log activation function.

The number of filters for the temporal and spatial convolution were fixed equal, and sampled from *f* ∼ *U*_N_[30, 50]. The length of the temporal filter was sampled from *l* ∼ *U*_ℕ_ [15, 35]. The number of strides for the mean pooling layer was sampled from *t* ∼ *U*_ℕ_ [10, 20], and the temporal length of the pooling layer was calculated as 5*t*. Finally, the drop out rate prior to the final layer was sampled uniformly between 0 and 0.5.

### 2.3. Domain adaptation

Two different methods for domain adaptation were implemented, (1) domain adversarial learning [29] and (2) CMMN [30]. While the former has received more attention in the field of DL, CMMN has the property that the Monge mapping convolutional filters can be computed at test time. We considered the scenario where no data from the test domain was available during training, hence domain adversarial learning can only be computed on the source data.

For LODO, an experiment used domain adversarial learning, CMMN, both, or no domain adaptation technique, with equal probabilities. For LODI, an experiment used either CMMN or no domain adaptation technique, sampled with equal probabilities.

#### 2.3.1. Domain adversarial learning with domain discriminator

Domain-adversarial learning using a gradient reversal layer was used as a method to obtain dataset invariant features [29]. For all DL architectures, it was implemented only for the last hidden layer, as this was the only vector representation available.

The optimisation with a domain discriminator includes adding a regularising term to the loss function, which penalises the extraction of domain specific features. To balance the regularisation term and the regression loss, a linear combination of the two is commonly used as

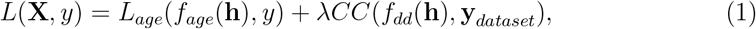

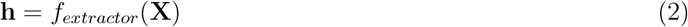

where **X** ∈ ℝ^*c*×*T*^ is the input EEG with channels and time steps as rows and columns, *y* ∈ ℝ is the target variable, *L*_*age*_(·, ·) is the loss used for the main regression task (either mean squared error or mean absolute error), *CC*(·, ·) is categorical crossentropy, *f*_*age*_ is the regression module of the DL model, *f*_*dd*_ is the domain discriminator, **y**_*dataset*_ is a one-hot encoded vector indicating which dataset the input data belongs to, *f*_*extractor*_ is the feature extractor of the DL model, and *λ* ∈ ℝ^+^ is a HP which determines the weighting of the discriminator loss. In the experiments, *λ* was sampled from the log uniform distribution *λ* = 10^*x*^, *x* ∼ *U* [−6, −1].

##### Discriminator architecture

The architecture of the discriminator was a multilayer perceptron, with number of hidden layers sampled from {0, 1, 2, 3, 4, 5} with equal probabilities. The size of the first layer was determined by uniform sampling from the distribution *D* = ⌊*x*⌋, *x* ∼ *U* [0.5*d*, 2*d*], where *d* ∈ N is the dimensionality of the input vector passed to the domain discriminator. The subsequent layers were decreasing exponentially in number of units by a factor sampled with equal probabilities from {2, 3}. Relu was used as the activation function for all hidden layers.

#### 2.3.2. CMMN

CMMN is a domain adaptation technique which provides test time adaptation to new datasets without the need for retraining the model [30]. It involves applying a Monge mapping convolutional filter per domain, which adapts the power spectral density (PSD) to a Wasserstein barycenter to obtain a homogeneous frequency spectrum.

In our study, the PSDs at subject level were estimated by averaging the PSDs computed per epoch. This was to avoid computing PSDs on discontinuous EEG data, as autoreject may discard data at epoch level. To compute the PSD barycenters, the grand average PSDs were first computed per channel and dataset. This was made to ensure equal representation of the datasets in the barycenter estimation. Then, we used Eq. 6 in [30] to compute the PSD barycenter of the *i*-th channel as

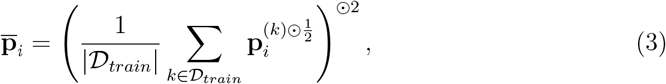

where *𝒟*_*train*_ denotes the datasets used for training, ·^⊙*n*^ denotes element-wise power of *n*, and 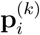 is the grand average PSD of the *i*-th channel of dataset *k*. The Monge mapping convolutional filters were computed per dataset and channel using Eq. 5 in [30] as

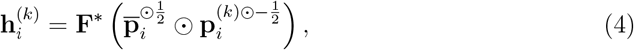

where **F**^∗^ is the inverse Fourier transform. Finally, the Monge mapping filter was applied per channel and dataset to the EEG signal prior to passing it do the DL model by (Eq. 5 in [30])

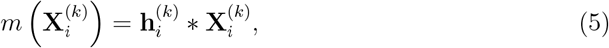

where 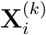 is the *i*-th channel of an EEG signal from the *k*-th dataset, and ∗ is the convolution operator.

Welch method was used to compute the PSDs, with the filter length sampled from the uniform distribution *U* [0.5*s*, 4*s*].

##### CMMN with RBP

As both PSD barycenters and Monge mapping convolutional filters were computed per channel, this required adaptation when the spatial dimension mismatch was handled using RBP rather than interpolation (see Sec. 2.5). We adopted a simple strategy, where PSD barycenters and Monge mapping convolutional filters were computed per region instead of per channel. The PSD of a region was computed on subject level as the average PSD of all channels within that region. Then, the grand average PSD of a region was computed as the mean of subject-level PSDs. The barycenter was followingly computed as in Eq. 3. The Monge mapping convolutional filters were applied after the RBP layer, on the region representations.

### 2.4. Sample re-weighting

Given the high variability of sample sizes across the datasets (see Sec. 2.1.1), a reasonable concern is that the largest datasets will dominate the loss during optimisation. This can be observed by reformulating the regression loss as

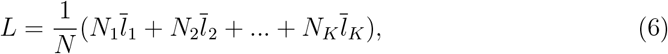

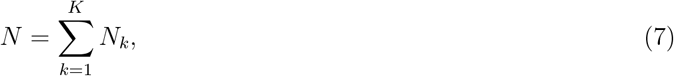

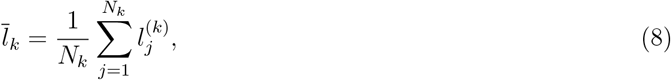

where *N*_*k*_ is the sample size of the *k*-th dataset, *K* is the number of datasets used for training, and 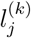 denotes the loss of the *j*-th subject from dataset *k*. As this may be an unwanted effect, a sample importance re-weighting was implemented. The importance re-weighting allows samples from smaller datasets to be weighted more, thus increasing the relative importance of the smaller datasets. The re-weighting was obtained by multiplying the loss computed per sample by a weight 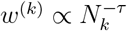, where *k* indexes the dataset, and *τ* ∈ [0, 1] is an HP which determines the power of up-weighting the smaller datasets. The two extremes correspond to equal sample importance (no re-weighting, *τ* = 0), and equal dataset importance (*τ* = 1). A normalisation constant *α* was used to obtain weighted averaging. This yielded the re-weighted loss to effectively be computed as

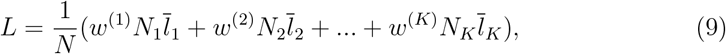

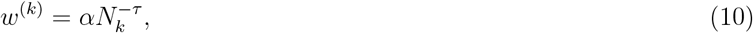

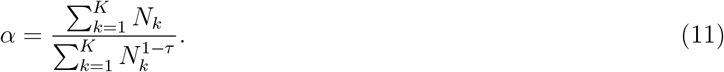

In our experiments, the sample re-weighting method was used with 50% probability. If the method was used, *τ* was sampled uniformly between 0 and 1. Furthermore, the normalisation constant *α* was computed using the sample sizes of the training datasets, not per batch.

### 2.5. Handling varied numbers of channels

Three different methods for handling a varied number of EEG channels were implemented. Those were Region based pooling (RBP) [44], spherical spline interpolation [45], and biophysical interpolation [46] with minimum-norm estimates (MNE). The selection of method for handling the varied electrode configurations in the different datasets was sampled with equal probability of using RBP and interpolation, and 50/50 for selecting spherical spline interpolation or MNE, if interpolation was used. For both interpolation methods, the fixed montage was sampled from those available in the training datasets with equal probabilities (see Sec. 2.1.1).

#### 2.5.1. HP distributions of RBP

##### Montage splitting

The algorithm for splitting the montages into distinct regions were as in the original work. All region formations were forced to be compatible with all datasets, which was obtained by enforcing compatibility with the channel system used in Miltiadous, as all other electrode configurations were essentially supersets of this. For each montage split, a split vector was sampled with equal probabilities from {**k**_1_, **k**_2_, **k**_3_, **k**_4_} with **k**_1_ = (2, 2, 2, 2, 2, 2, 2)^T^, **k**_2_ = (3, 3, 3, 3, 3, 3, 3, 3)^T^, **k**_3_ = (2, 3, 2, 3, 2, 3, 2, 3, 2)^T^, **k**_4_ = (4, 3, 2, 3, 4, 3, 2, 3, 4)^T^. A stopping criterion which prevents the split algorithm from creating new regions if the number of channels in a region is less than allowed was employed. The minimum number of allowed channels in a region was sampled uniformly from *U*_ℕ_ [1, 5] per montage split. The number of montage splits was sampled from a log uniform distribution *n* = [2^*x*^], *x* ∼ *U* [0, 4].

##### Pooling methods and modules

While the original work introduced four different pooling methods, we only considered three of them: averaging, channel attention, and channel attention with a head region. Continuous channel attention was not considered due to high memory and time consumption, with no performance gain. Furthermore, multiple pooling methods was allowed to be used in parallel, to increase the heterogeneity of the region representations. Parameter sharing between similar pooling methods of the different montage splits was allowed, and full parameter sharing between the montage splits was used with 50% probability. If full parameter sharing was not used, the number of *pooling modules* was sampled from *k* = *max*(*x*, number of montage splits), *x* ∼ *U*_N_[1, 10]. A single pooling module uses parameter sharing across a subset of the montage splits with the same pooling method. After sampling the number of pooling modules, the number of montage splits for the pooling modules were assigned by generating *k* bins with sizes equal to one, and iteratively incrementing the size of a randomly selected bin *n* − *k* times. The *i*-th bin size determined the number of montage splits for the *i*-th pooling module. For pooling modules using channel attention or channel attention with a head region, the number of convolutional kernels as sampled in ROCKET [47], was sampled from a log uniform distribution *n*_*kernels*_ = [10^*x*^], *x* ∼ *U* [2, 3], while the maximum receptive field was sampled from the log uniform distribution max receptive field = [10^*x*^], *x* ∼ *U* [2, 2.5]. For pooling modules using channel attention with a head region, the dimensionality of the search vector was sampled from *dim*(search vector) = [2^*x*^], *x* ∼ *U* [3, 7]. Finally, the weights of the head region and non-head regions were shared with a 50% probability. That is, the parameters of the embedding functions used in Eqs. (3-4) and (6-7) in [44] were either shared or not shared, sampled with equal probabilities.

### 2.6 Training HPs

All models were run for 100 epochs, and the batch size was fixed to 128. The Adam [31] optimiser was used for all experiments, which HPs were sampled as described in table 1. The regression loss function was either mean squared error (MSE) or mean absolute error (MAE), sampled with equal probabilities. After splitting into source and target datasets for a single fold in cross validation, a validation set was separated from the training set. This validation set consisted of 20% of the source data. In LODO, we ensured that 20% was used for validation per dataset.

**Table 1.**
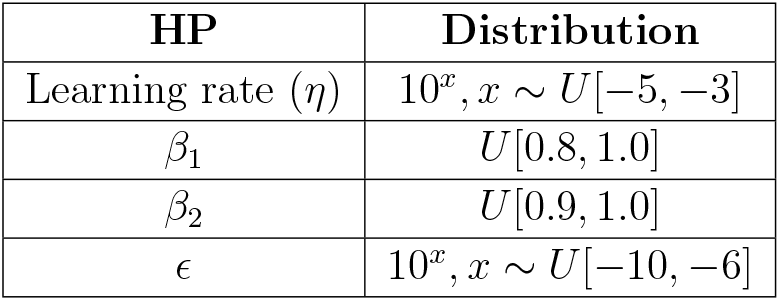
Sampling distributions for the Adam optimiser.

### 2.7. Experiments

Initially, the experiments ran for approximately 2 months, giving a total of 1087 initiated experiments. After these two months of experiments, an error in one of the HP distributions was discovered, as normalisation was always applied if RBP was used as the method for handling the varied electrode configurations. To balance this, we initiated 1087 new experiments with the exact same distributions, with the exception that normalisation was not applied if RBP was used. In total, 1110 LODO and 1064 LODI experiments were initiated. Mainly due to hardware limitations, 191 (LODO) and 178 (LODI) of these were not finalised, resulting in 919 and 886 experiments suitable for analysis for LODO and LODI, respectively. An overview of the HPs of the experiments which did not finalise is shown in figure A1.

The experiments were conducted on a computer equipped with an NVIDIA GeForce RTX 3060 12GB GPU.

## 3. Results

### 3.1. Performance estimates

Figure 3 shows the performance scores for all pairs of source and target datasets. When “Pooled” is used as the source dataset, it means that all datasets, except the specified target dataset, are used for training. Similarly, when “Pooled” is used as the target dataset, it refers to considering all datasets, except the source dataset, as a single combined dataset. 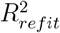 refers to the *R*^2^ score after fitting an intercept to the target dataset to ensure that the average prediction is equal to the average age [48]. Therefore, this metric requires a leakage of the average age of the target dataset. For each entry, the model was selected by maximising the performance on the validation set, to obtain unbiased performance estimates. The metric used for model selection was the same as the target metric for each of the heatmaps, with the exception of 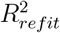, which used *R*^2^ for model selection. In Figures A2 (LODO) and A3 (LODI), model predictions are shown for each source/target dataset pair.

**Figure 3.**
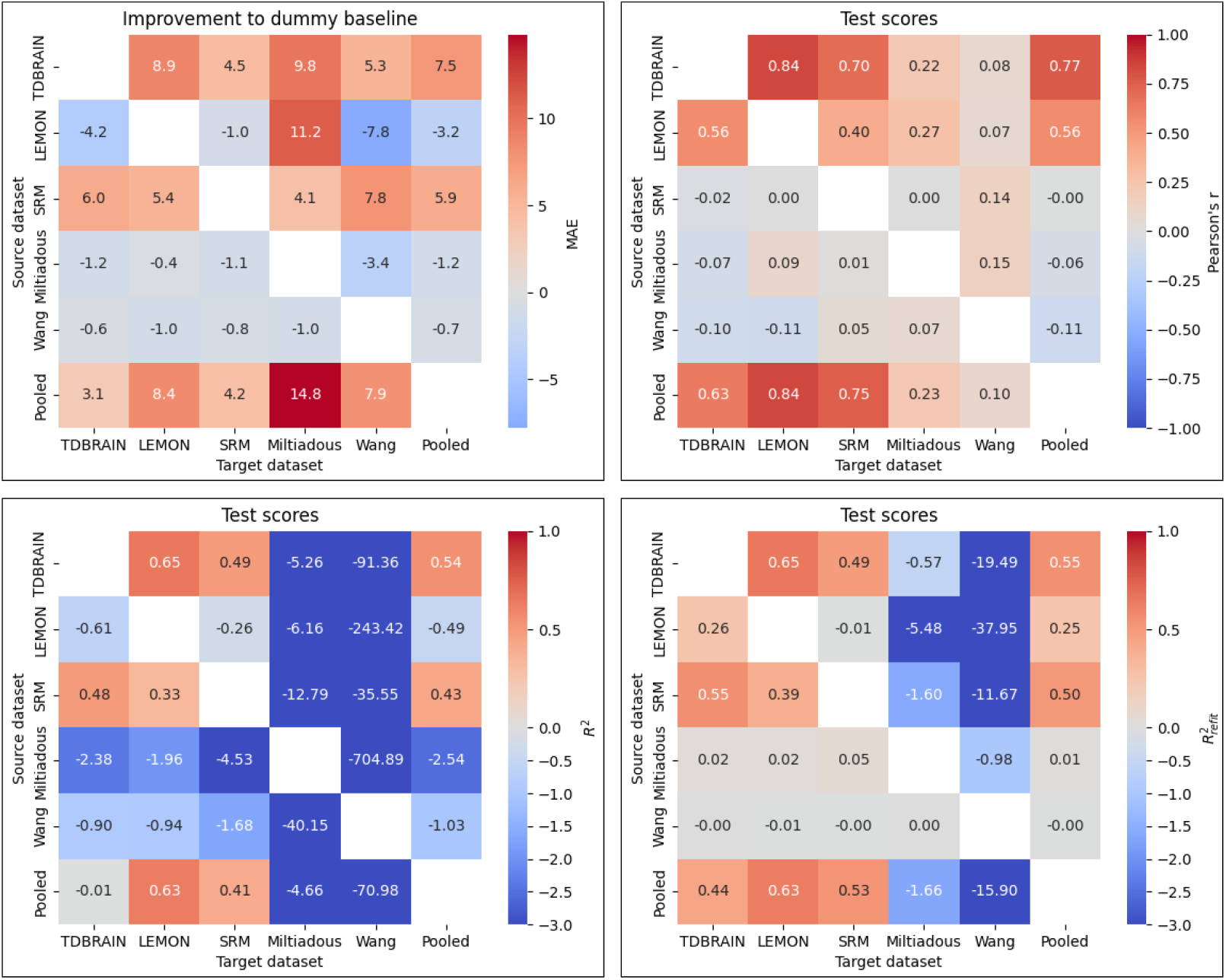
Performance scores for all pairs of source/target datasets show considerable variations depending on the source/target pair and the performance metric. For MAE, the scores reflect improvement over a dummy model that always predicts the average age of the source dataset, with a positive score indicating an improvement over this baseline and a negative number indicating a worsening from this baseline. For *R*^2^, negative scores indicate worse performance than guessing the average age of the target dataset. Consistent MAE improvements to baseline are observed only when TDBRAIN, SRM, or Pooled are used as source datasets. The correlations are highest for TDBRAIN, LEMON, and Pooled, which are the datasets with the highest sample sizes and the widest age distributions. Furthermore, as evident in figure 5, the poor Pearson’r scores with SRM as the source dataset are likely due to poor model selection. The *R*^2^ scores are particularly poor on the Miltiadous and Wang datasets, which are the smallest datasets with the most narrow age distribution. Refitting the intercept to ensure zero average error on the target dataset generally improved the performance score.

### 3.2. Performance distributions

Figures 4 and 5 show the distribution of performance scores for the LODO and LODI experiments, respectively, for different band-pass filters and DL architectures. For LODI, we selected the performance score from a single source dataset to the pooled target dataset. For each HPC, the epoch was selected by maximising the performance of the respective metric on the validation set. In case the prediction array was constant, Pearson’s r was set equal to zero.

**Figure 4.**
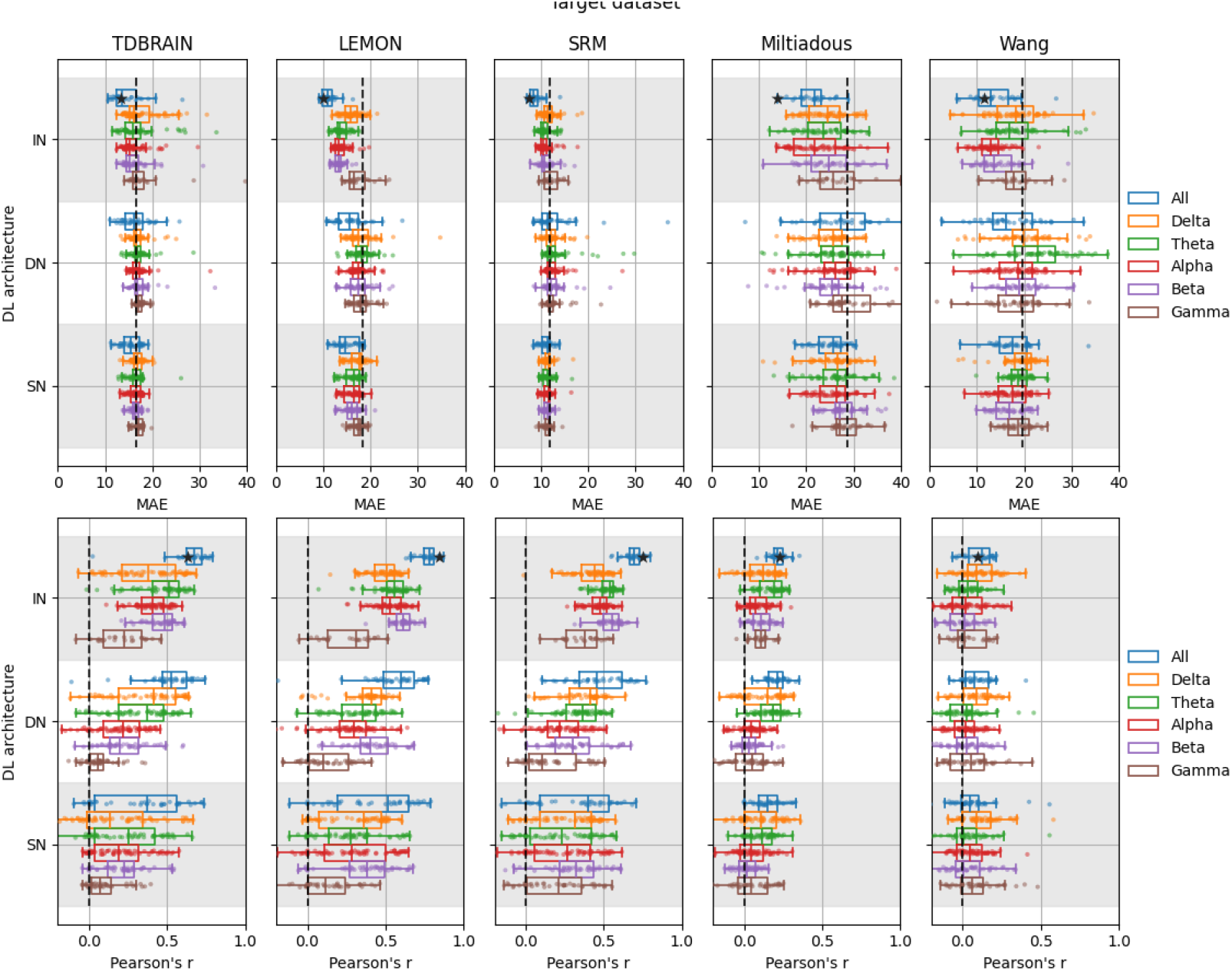
The distribution of performance scores across all LODO experiments, for different DL architectures and band-pass filters. The position of the star indicates the DL architecture, band-pass filter, and performance score after model selection, and the dotted line indicates the performance of a dummy model which always guesses the average age of the source dataset. The use of Inception network and considering the frequency range from 1 to 45 Hz was a particularly strong HP combination. Abbreviations: IN (Inception network), DN (Deep4Net), SN (ShallowFBCSPNet).

**Figure 5.**
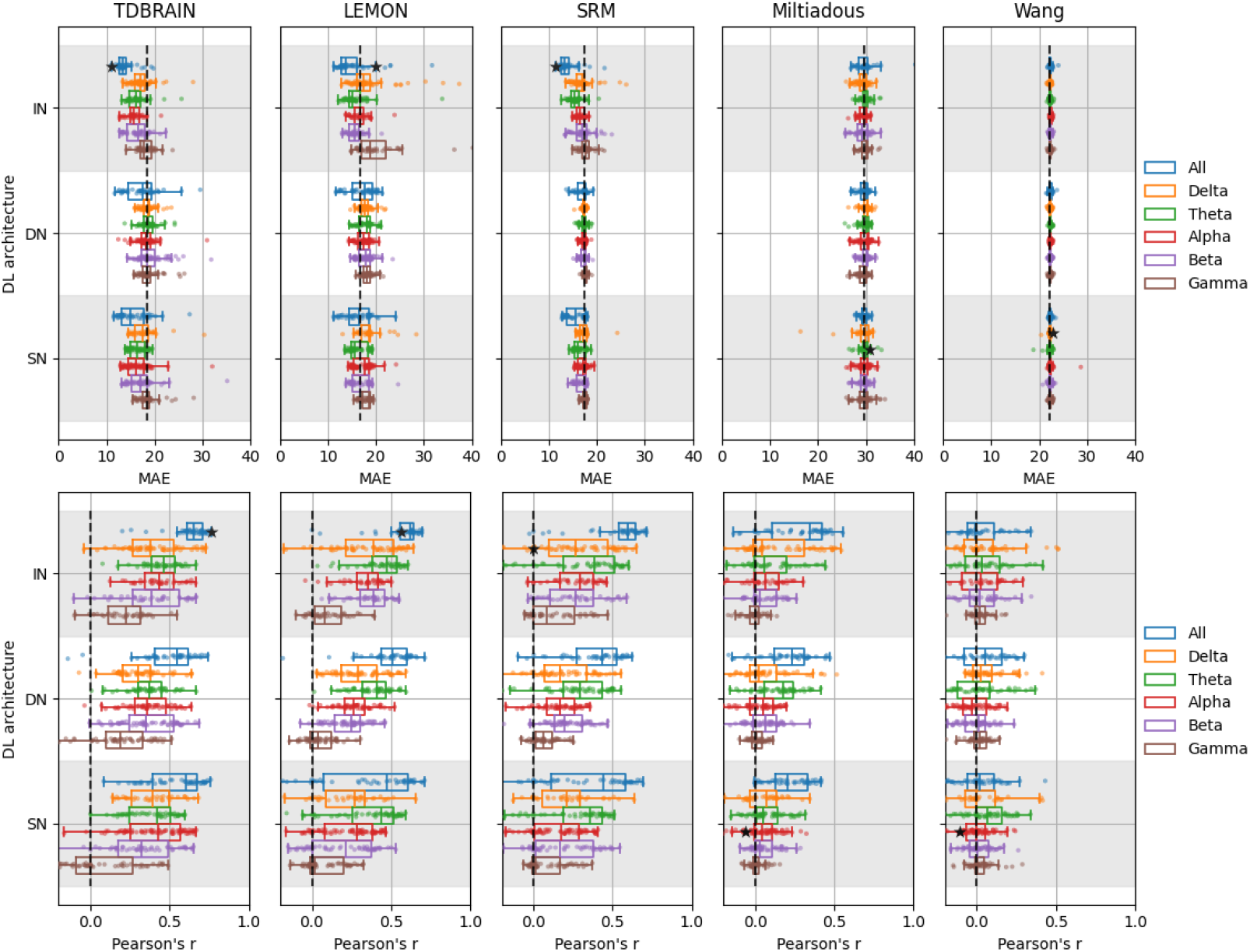
The distribution of performance scores across all LODI experiments when pooling the target datasets, for different DL architectures and band-pass filters. The position of the star indicates the DL architecture, band-pass filter, and performance score after model selection, and the dotted line indicates the performance of a dummy model which always guesses the average age in the source dataset. The use of Inception network and considering the frequency range from 1 to 45 Hz was a particularly strong HP combination. Furthermore, it is evident that model selection occasionally failed to identify a high-performing model. This is exemplified by a Pearson’s r score that was no better than the dummy model when the SRM or Miltiadous dataset was used as the source dataset, despite most models employing a band-pass filter between 1 and 45 Hz outperformed this baseline. Abbreviations: IN (Inception network), DN (Deep4Net), SN (ShallowFBCSPNet).

### 3.3. HP importance

Figure 6 shows the HP importance scores computed using the fANOVA method [49], in the region of configuration space which represents a score greater than the 95th percentile. This method fits a random forest model to predict the performance given the HPCs, performs functional decomposition, and computes the variance of the functional components per subset of HPs. In the figure, we show a boxplot of these variances across 64 trees, all trained with a maximum depth of 64. We selected Pearson’s r as the performance and selection metric and applied a lower bound at the 95th percentile. This ensured that all performance variations represent an improvement relative to this high performance score [49].

**Figure 6.**
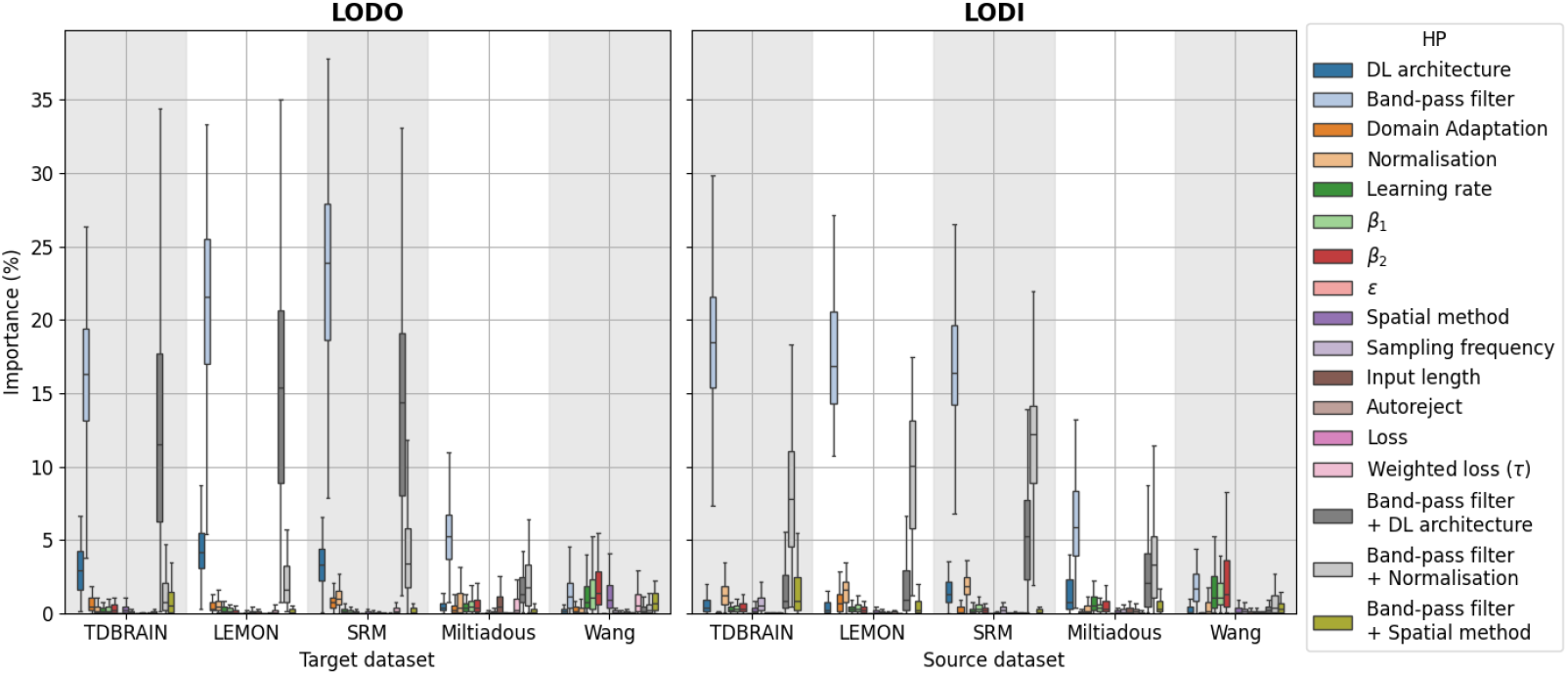
Hyperparameter importance as computed by fANOVA [49], for explaining performance variation above the 95th percentile. This is done by fitting a random forest model to predict the performance based on hyperparameter configurations (HPCs), and then decomposing the performance variations above the 95th percentile into contributions from subsets of hyperparameters. The marginal importance of band-pass filters is particularly high across most datasets for both LODO and LODI. For LODO, an interaction effect between band-pass filter and DL architecture is observed for TDBRAIN, LEMON, and SRM. The most important interaction effect for LODI was band-pass filter and normalisation, followed by interaction effects between band-pass filter and DL architecture. Finally, the effective HP dimensionality was considerably lower than the total number of HPs, as the performance variations were attributable to only a small subset of them.

## 4. Discussion

This study investigated the ability of DL based brain age prediction models to generalise to fully unseen and external EEG datasets.

EEG may serve as a convenient modality for AI applications due to its availability and low acquisition and use cost compared to other imaging techniques. Our research question examines the robustness of EEG-based DL models, whose clinical utility is highly dependent on their ability to work across diverse datasets, populations, and acquisition hardware. Model generalisation is particularly important for problems where data on both the EEG and the response variable exist at only one or a few clinical sites. In that setting, even supervised model configuration can be unfeasible, as the majority of clinical sites lack data on the response variable. For the dementia risk assessment scenario, which is the objective of the AI-Mind study [3], several years of data collection is necessary. In this case, the EEG should be collected in the MCI stage, and therefore, the clinical outcome may remain unknown for years after the initial measurement.

### 4.1. Main findings

In sum, our results indicate that brain-age related characteristics extracted by DL models can have some transferability to unseen datasets. This is reflected by strong correlation coefficients in three of the datasets for the leave-one-dataset-out (LODO) experiments, and similar results for the same three datasets in the leave-one-dataset-in (LODI) experiments. For the remaining two datasets, the age ranges were narrower (see Figure 2), which may explain the poorer performance. Additionally, the Miltiadous dataset contains participants with frontotemporal dementia and Alzheimer’s disease. As such neurodegenerative diseases are associated with various alterations of the EEG, this could potentially further extend the dataset beyond the training distribution.

For comparison, brain age prediction from M/EEG data was benchmarked in [27], finding *R*^2^ scores between 0.60 and 0.74, which is comparable to our best results (see Fig 3). Most comparably, they reported an *R*^2^ = 0.69 using Shallow and DeepNet on the LEMON dataset. While those scores are better than those in the current study, our results were obtained without including any data from the LEMON dataset for training. It is therefore not expected to obtain similar performance to studies which have not separated the entire dataset from training. This is highly relevant for metrics which are not location and scale invariant, as the DL models are likely to be biased towards the mean and scale of the age distribution in the training set. A previous study has also reported *R*^2^ = 0.81, *MAE* = 5.96 and Pearson’s r = 0.9 on the TDBRAIN dataset, although they also included EEG data recorded with the eyes open [50]. While the novelty in our results is the demonstration of cross-dataset generalisation, it is evident that our models did not reach state-of-the art performance on brain age prediction as a regression task. However, factors other than model generalisation may be the cause of this, such as (1) they used different pipelines (different architecture, they included both eyes open and eyes closed, and they applied data augmentation), (2) our testing procedures were different (we tested on the entire TDBRAIN dataset, whereas they used 10-fold cross validation), and (3) by not considering any data from the TDBRAIN dataset, the sample size used for training decreases substantially.

It is a common concern that DL models will fail when applied to new datasets, with different acquisition hardware and a different population. Although our work has limitations, as discussed in Sec. 4.4, our results demonstrate promising cross-dataset generalisation of DL models on resting-state EEG data. Therefore, we encourage future studies to use external datasets for testing such models, when possible.

We emphasize that our results are unlikely to be transferable to other neuroimaging modalities, with the possible exception of magnetoencephalography (MEG). This is because imaging techniques such as MRI, fMRI, and PET contain features that are highly different from EEG, seen from a signal processing perspective. While images contain features such as edges and texture, resting-state EEG contains oscillations with prominent frequency bands, and is better characterised by amplitude and phase information.

A further consideration in the context of brain age models is that they are usually developed for constructing biomarkers of healthy and pathological brain aging. Maximising the performance of such models is not equivalent to maximising their clinical utility [51, 52]. For example, one study [53] demonstrated that MRI-based brain age models can benefit from being moderately fit for the purpose of pathology discrimination. Similarly and importantly, our models not reaching state-of-the-art performance on regression metrics does not necessarily degrade their potential clinical utility.

Our work further demonstrates that the model selection procedure can exhibit high variance, in particular when the sample size of the validation set is small. This was seen in figure 5, where the performance of the selected model on the pooled test dataset after training on SRM or Miltiadous was equal to or worse than the dummy baseline, despite the majority of models employing a band-pass filter between 1 and 45Hz outperformed this baseline. These suboptimal scores likely occurred because fluctuations from random variations are more pronounced with small sample sizes. Although this is an unsurprising finding, it empirically underscores the difficulties in model selection for low sample size regimes.

### 4.2. Related work

A similar study was made on the inter-site generalisability of a brain age prediction model on preterm infants [54] with two different EEG datasets from different sites (*N*_1_ = 37 and *N*_2_ = 36). A brain age prediction model using a set of 43 hand-crafted features was developed. They found that the performance on the test dataset was improved when restricting the feature set to those with no statistically significant difference between the two datasets, which may be considered unsupervised feature extraction. This procedure, however, may not always be feasible in practice, as it requires access to data from the target clinical site during the time of model training. They further highlight that EEG based classifiers are challenged by site specific differences.

Another similar study developed an interpretable machine learning model to predict age from sleep EEG data [55]. To investigate the impact of the training data, they trained the model on 2365 EEGs from healthy subjects from the MGH dataset and tested the model on 1974 paired EEGs from two different visits of the SHHS dataset. They obtained Pearson’s correlation coefficients of 0.61 and 0.56 on visit 1 and 2, respectively, which is similar to our results.

In [56], quantitative EEG (qEEG) norms from cross-spectral matrices (HarMN-qEEG) were calculated using data from 9 countries, 12 devices, and 14 studies, with a total of 1564 subjects. The study identified qEEG batch effects across different sites and demonstrated that employing harmonized Riemannian norms improved alignment and diagnostic accuracy. Using both simulated and the HarMNqEEG data, [48] showed the benefit of geodesic optimisation for predictive shift adaptation (GOPSA) when adapting to data from unseen sources, achieving *R*^2^ scores as high as 0.8 on HarMNqEEG. Similar to our results, however, the performance scores varied depending on the source/target pairs.

Brain age predictions across heterogeneous datasets has been studied using MRI data. [52] found that the most limiting factor of generalisation was acquisition protocol differences and biased brain age estimates. [24] obtained reasonable generalisation to an independent dataset acquired at a different site. By training on a dataset with variations in geographic locations, scanners, acquisition protocols, and studies, [53] obtained robust brain age estimates without the need for specialised preprocessing on data from an unseen site.

### 4.3. Impact of HPs

Interestingly, using all frequency bands was one of the most important design choices for both the LODO and LODI experiments. This is evident as band-pass filtering was the most important HP for the high performing regions of the configuration space in both LODO and LODI. A likely explanation is that band-pass filtering to a single frequency band removes an excessive amount of predictive information. This is in line with a recent study observing that for age-prediction on the TDBRAIN dataset using Riemannian models, the combination of all frequencies led to substantially better performance compared to any individual frequency or their average [28]. Although it is theoretically possible that the DL models have some bias which makes the generalisation on single frequency bands poor, applying additional pre-processing steps should lead to more homogenised datasets. If the age predictive information was restricted to a single frequency band, we argue that it should be easier to obtain good results using that frequency band only. A recent study tested this hypothesis more directly using learnable wavelets and found that, for age prediction, multiple frequency filters were required, while on other prediction tasks 2-3 filters were sufficient for optimal performance [57].

Although our study was intended mainly to investigate the behavior of DL-based prediction models on resting-state EEG data when applied to unseen datasets, this finding adds to the growing literature on brain-age modeling with EEG. [58] observed age-related changes in the alpha frequency band for two different datasets, containing participants aged 5-18 years (*N*_1_ = 27 and *N*_2_ = 86). [59] obtained highest age predictive value from the beta band, using a deep BLSTM-LSTM architecture. [25] found significant differences between old and young subjects in delta sources in the occipital area. Furthermore, low and high alpha was found to have significantly lower power in parietal, occipital, temporal, and limbic areas in older subjects. Further analysis suggested both linear and non-linear trends of occipital delta and posterior cortical alpha rhythms with aging. Building on our finding that the best predictions came from considering all frequency bands, our study emphasises the importance of considering the entire EEG for predictive brain age modelling, rather than restricting the analysis to features and characteristics of a single frequency band.

The second most important HP effect was dependent on the experimental setup. For LODO, the interaction effect between band-pass filter and DL architecture was the most important, whereas for LODI, the interaction effect between band-pass filter and normalisation was the most prominent. For LODI, and depending on the source dataset, there were varied interaction effects between band-pass filter and DL architecture. This highlights the importance of HP optimisation, as different experimental setups and different datasets may exhibit variation in optimal HPCs and their importance. That is, transferability in successful HPCs from our results to a next similar scenario is not guaranteed, and therefore, we recommend that future studies employ comprehensive HP optimisation for their specific scenario.

The impact of processing with autoreject [38] did not stand out as a factor affecting cross-dataset generalisation. Recent work pointed out that denoising with autoreject could improve prediction performance within datasets for age- and sex prediction[28]. It is noteworthy, however, that the reported effect was more pronounced for weaker, non-deep prediction models, the results referred to within-dataset performance with high baseline performance, and no other normalisation or domain-adaptation steps were applied. It is conceivable that the complex DL architectures, in combination, with normalisation steps performed implicit denoising and that this effect may be more pronounced once a certain performance level is reached. We, therefore, recommend continued attention to the role of EEG preprocessing in the context of machine learning for EEG.

### 4.4. Limitations

For all experiments performed in this study, EEG was represented as multivariate time series. Other representations which can be used include time-frequency domain as well as using the power spectra. Furthermore, we limited the experiments to using eyes-closed resting state data only. Combining data from both eyes open and eyes closed data gave improved results in [50]. However, we hypothesised that training and testing on more homogeneous data would serve as a better starting point, given the current lack of similar studies directly focusing on cross-dataset generalisation using DL on EEG data.

Only a single dataset was collected from a non-European country. This dataset from [37], however, has a narrow age range, making it unfeasible to properly test brain age prediction models (see figure 2), in particular as a source dataset.

Although this study documented an extensive set of experiments with various HPs and design choices, the results are unlikely to reflect those achievable with a globally optimal configuration. New architectures, techniques, and different configurations may potentially improve the results. This includes pipelines for pre-processing and feature extraction, such as variations in filter construction hyperparameters. Furthermore, this study investigated only LODO and LODI cross validation. It could, however, be beneficial to train on other subsets of the available datasets. As the brain age models were most successful on the three datasets TDBRAIN, LEMON, and SRM, an interesting approach may be to only use those datasets for training and testing.

Finally, as the primary aim of this study was to investigate the generalisation robustness of DL models on EEG data, associations of brain age with pathological conditions were not investigated. Furthermore, since brain age as a pathological biomarker has been limited to MRI data, the clinical utility of our resulting DL models remains unvalidated.

## 5. Conclusion

This study investigated the ability of deep learning models trained on EEG data to generalise to new and unseen datasets, with variations in populations and acquisition hardware. A broad range of hyperparameter configurations were tested to improve the reliability of the results, and yield robust benchmarks. Our results, although varying with respect to the target and source datasets, demonstrated systematic correlations between predicted age and chronological age on unseen datasets with the broadest age distributions. Poor model selection was observed in the low sample size regime. Using all frequency bands was the most important design choice for obtaining high prediction scores, and the second most important hyperparameter effect depended on the experimental setup, which highlights the importance of comprehensive hyperparameter optimisation. In sum, our study provides a set of benchmarks for assessing and further developing machine-learning methods that tackle cross-dataset learning from EEG signals.

## Acknowledgments

This project has received funding from the European Union’s Horizon 2020 research and innovation programme under grant agreement No 964220. This publication reflects views of the authors and the European Commission is not responsible for any use that may be made of the information it contains.

DE is a full-time employee of F. Hoffmann-La Roche Ltd.

## Author contributions

**Thomas Tveitstøl**: Conceptualization, Data curation, Formal analysis, Investigation, Methodology, Software, Visualization, Writing – original draft **Mats Tveter:** Visualization, Writing – review & editing **Christoffer Hatlestad-Hall:** Writing – review & editing **Hugo L Hammer:** Supervision, Writing – review & editing **Denis A Engemann:** Formal analysis, Supervision, Writing – review & editing **Ira R J Hebold Haraldsen:** Funding acquisition, Supervision, Writing – review & editing

## Appendix

### Appendix A.1. Failed models

**Figure A1.**
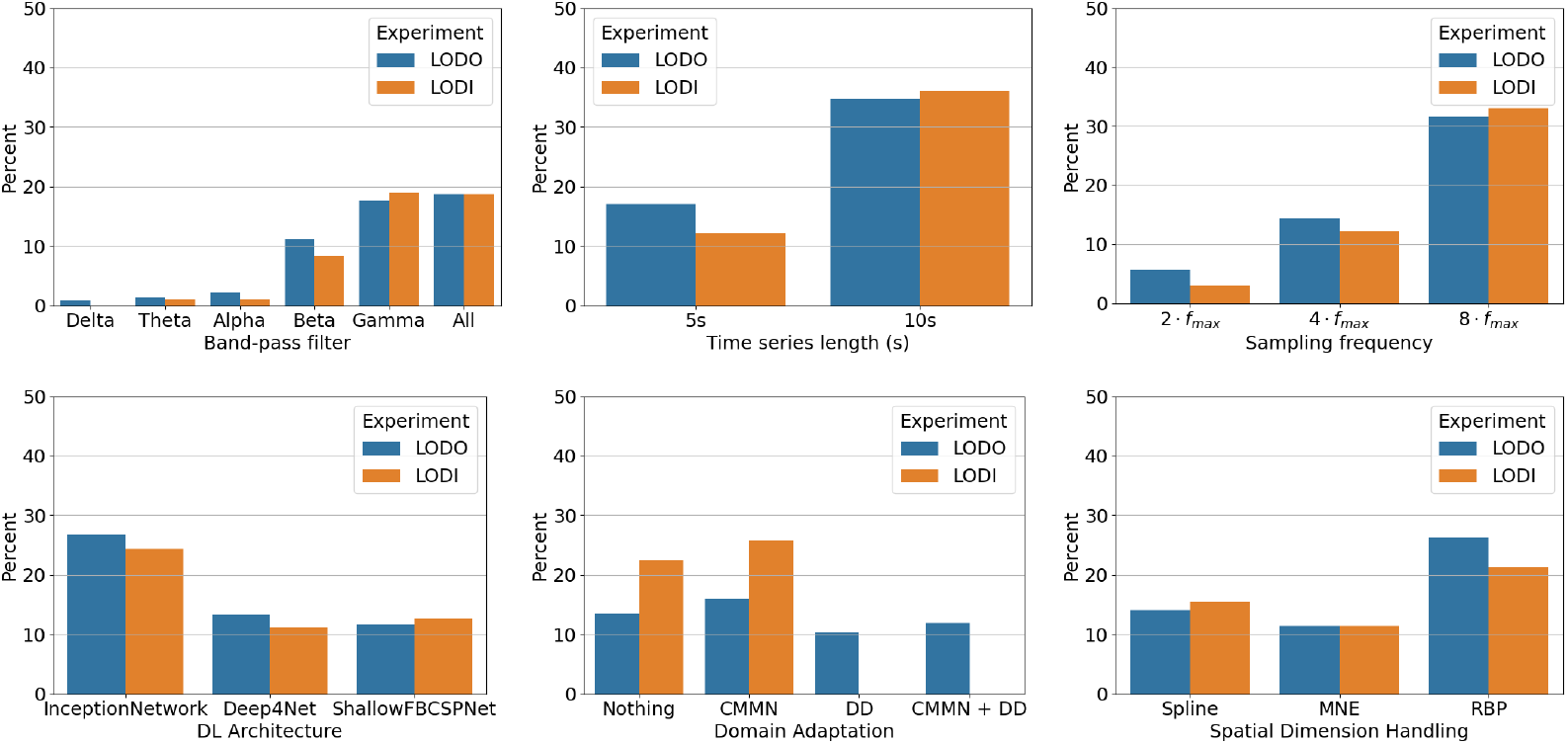
HP distributions of the models which did not finish successfully.

#### Appendix A.2. Prediction plots

Figures A2 (LODO) and A3 (LODI) show scatterplots of the model predictions per source and target pair, for different metrics in model selection.

#### Appendix A.3. Test performance vs validation performance

Figures A4 (LODO) A5 (LODI) show scatterplots of test performance as a function of validation performance per source and target pair.

**Figure A2.**
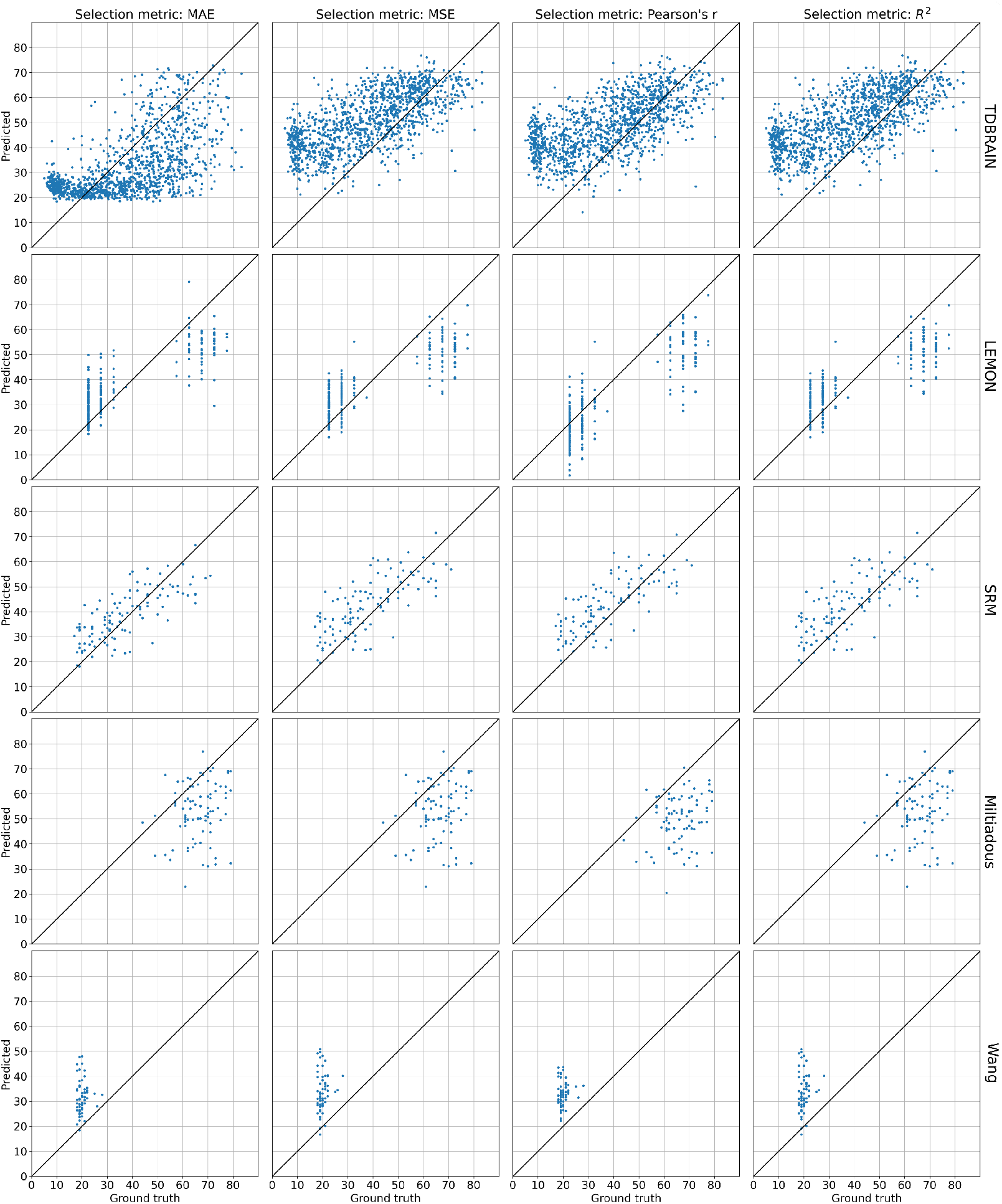
Test predictions of the selected models for all source/target dataset pairs in LODO, using different selection metrics.

**Figure A3.**
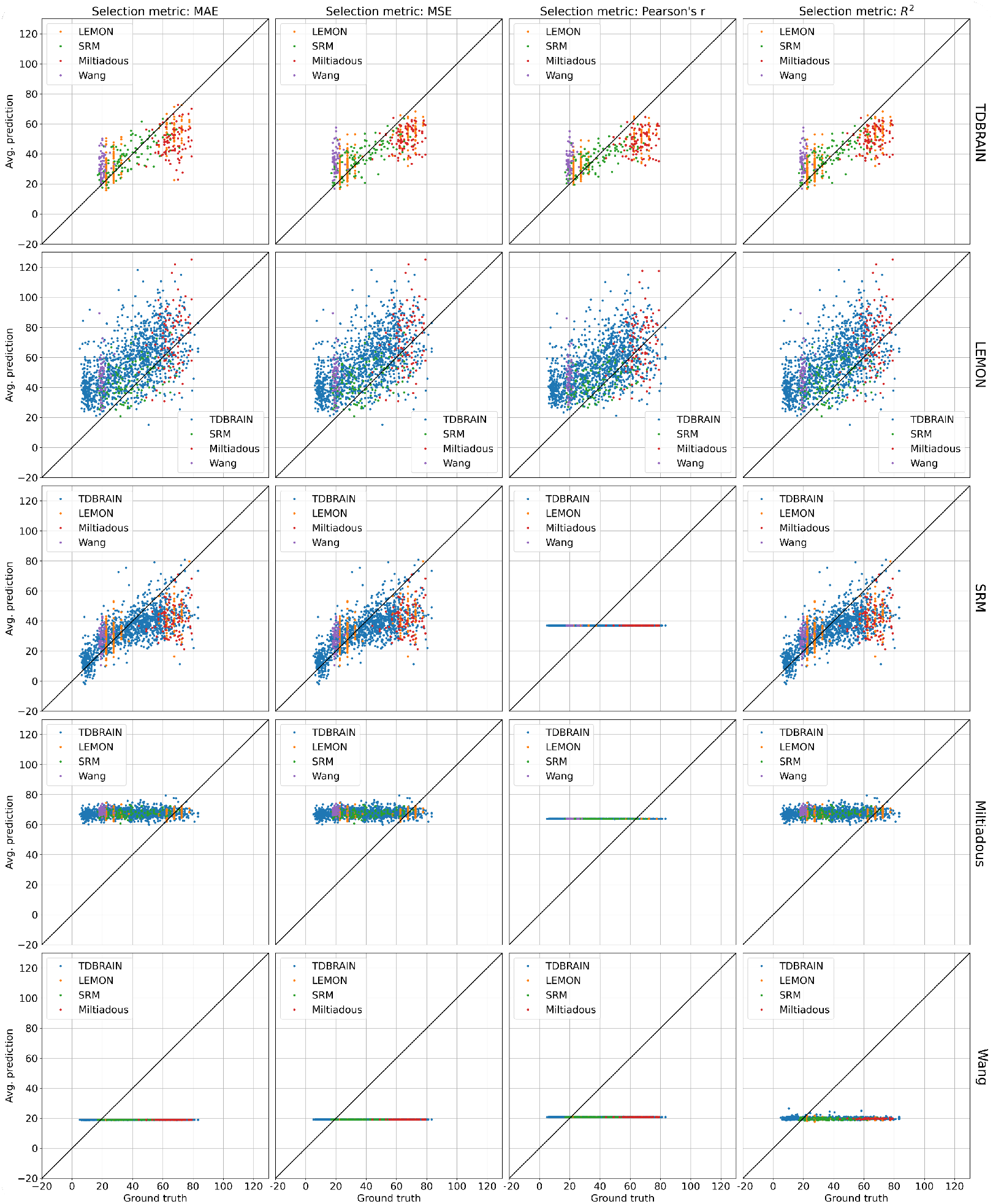
Test predictions of the selected models for all source/target dataset pairs in LODI, using different selection metrics.

**Figure A4.**
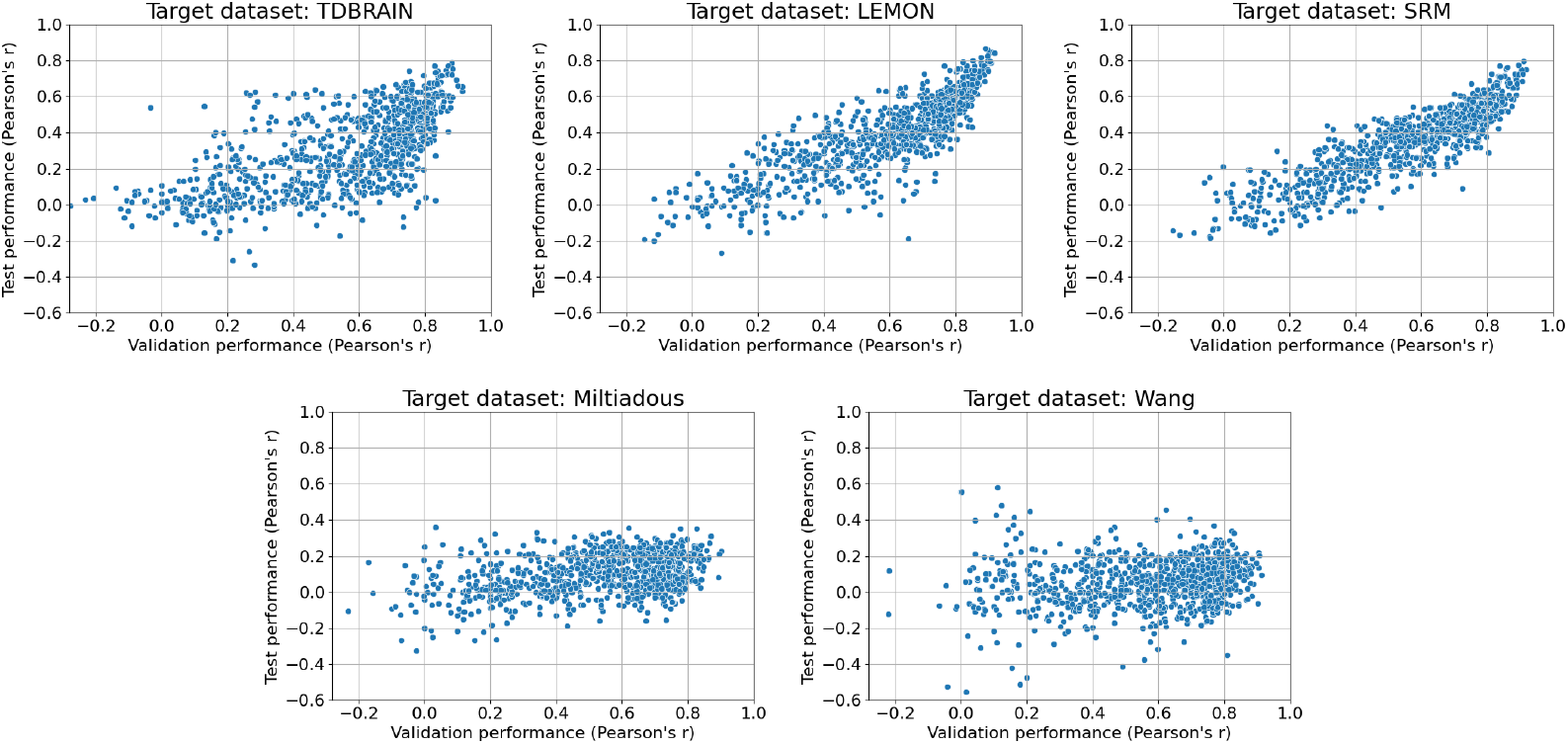
Test performance plotted against validation performance on the pooled dataset, for the different target datasets using leave-one-dataset-out cross validation.

**Figure A5.**
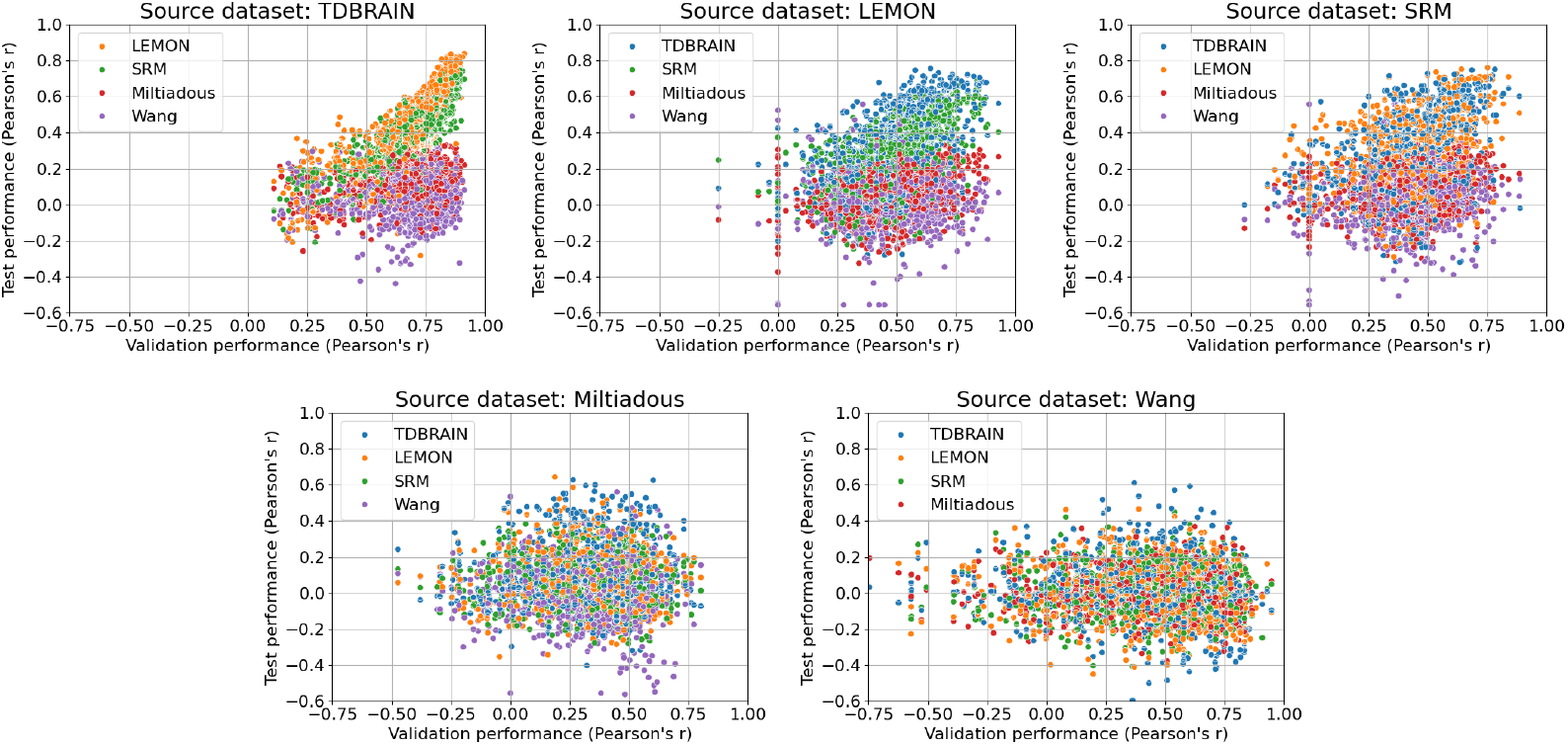
Test performance plotted against validation performance for the different datasets using leave-one-dataset-in cross validation.

